# Heterogeneous CaMKII-dependent synaptic compensations in CA1 pyramidal neurons from acute slices with dissected CA3

**DOI:** 10.1101/2021.11.09.467941

**Authors:** Pablo Vergara, Gabriela Pino, Jorge Vera, Magdalena Sanhueza

## Abstract

Prolonged changes in neural activity trigger homeostatic synaptic plasticity (HSP) allowing neuronal networks to operate in functional ranges. Cell-wide or input-specific adaptations can be induced by pharmacological or genetic manipulations of activity, and by sensory deprivation. Reactive functional changes caused by deafferentation may partially share mechanisms with HSP. Acute hippocampal slices constitute a suitable model to investigate relatively rapid (hours) pathway-specific modifications occurring after denervation and explore the underlying mechanisms. As Schaffer collaterals constitute a major glutamatergic input to CA1 pyramidal neurons, we conducted whole-cell recordings of miniature excitatory postsynaptic currents (mEPSCs) to evaluate changes over 12 hours after slice preparation and CA3 dissection. We observed an increment in mEPSCs amplitude and a decrease in decay time, suggesting synaptic AMPA receptor upregulation and subunit content modifications. Sorting mEPSC by rise time, a correlate of synapse location along dendrites, revealed amplitude raises at two separate domains. A specific frequency increase was observed in the same domains and was accompanied by a global, unspecific raise. Amplitude and frequency increments were lower at sites initially more active, consistent with local compensatory processes. Transient preincubation with a specific Ca^2+^/calmodulin-dependent kinase II (CaMKII) inhibitor either blocked or occluded amplitude and frequency upregulation in different synapse populations. Results are consistent with the concurrent development of different known CaMKII-dependent HSP processes. Our observations support that deafferentation causes rapid and diverse compensations resembling classical slow forms of adaptation to inactivity. These results may contribute to understand fast-developing homeostatic or pathological events after brain injury.

## INTRODUCTION

Homeostatic synaptic plasticity (HSP) are compensatory mechanisms adjusting synaptic strength up or down depending on activity levels and maintaining average transmission within a range allowing circuit stability (Turrigiano and Nelson, 2000). Inactivity-induced upregulation of glutamatergic synapses is expressed after long-lasting (typically, 1-2 days) pharmacological inhibition of action potentials or synaptic transmission (G Turrigiano et al., 1998; O’Brien et al., 1998), and in cortical slices from sensory-deprived rodents (Goel et al., 2006; Lee and Kirkwood, 2019).

While pioneer works described a drug-induced cell-wide synaptic scaling, locally regulated homeostatic processes have now been extensively reported (Turrigiano, 2012) Diverse expression mechanisms have been described for adaptation to inactivity. In many cases, they include the specific synaptic incorporation of GluA2-lacking, Ca^2+^-permeable AMPA receptors (Thiagarajan et al., 2005; Sutton et al., 2006). In others, both GluA1 and GluA2 subunits are upregulated (O’Brien et al., 1998; Wierenga, 2005).

Ca^2+^/calmodulin-dependent kinase II (CaMKII) is a major component of glutamatergic synapses and is crucial for Hebbian plasticity (Lisman et al., 2012; Hell, 2014; Bayer and Schulman, 2019). In the forebrain, CaMKII holoenzyme is usually formed by 12 subunits of the α and β isoforms (Bennett et al., 1983; Lisman et al., 2012). Both Ca^2+^-dependent and autonomous CaMKIIα activity (generated by T286 autophosphorylation), and its binding to the glutamate NMDA receptor (NMDAR) are critical for long-term potentiation (LTP). Moreover, the CaMKIIα/NMDAR interaction is involved in synaptic strength maintenance (Sanhueza et al., 2011; Incontro et al., 2018). CaMKII also participates in forms of HSP (Lee, 2012; Hell, 2014). α and β subunits have different affinities for Ca^2+^/CaM and interact with specific cytoskeletal and postsynaptic density proteins, determining holoenzyme localization in an activity-dependent manner (Hell, 2014). Subunit expression is inversely regulated by neural activity: long-lasting inactivity increases CAMKIIß and decreases CaMKIIα. Moreover, CAMKIIß knockdown or inhibition by the CaMKII inhibitor KN-93 prevents the synaptic upregulation caused by activity blockade (Thiagarajan et al., 2002; Groth et al., 2011). On the other hand, CaMKIIα normally limits AMPAR synaptic incorporation and network silencing disrupts this mechanism, leading to homeostatic upregulation (Wang et al., 2011; Djakovic et al., 2012).

*In vivo* studies have shown that denervation induces compensatory adaptations in peripheral and central synapses, days after deafferentation (Li and Prince, 2002; Houweling et al., 2005; Turrigiano, 2012). Moreover, surgical removal of the entorhinal cortex from slice cultures triggers pathway-specific mEPSC amplitude upregulation in *dentate gyrus* granule cells; changes take ~2 days and share signaling properties with forms of drug-induced HSP (Vlachos et al., 2012; Perederiy and Westbrook, 2013; Bissen et al., 2021). In addition to contributing to traumatic nerve injury studies, deafferentation models could help to resolve input-dependent homeostatic compensations. Long-lasting action potential inhibition by tetrodotoxin (TTX) in the hippocampus *in vivo* or in slice cultures induces complex cell-type-dependent synaptic adaptations (Echegoyen et al., 2007)(Kim and Tsien, 2008). As the whole circuit is altered, it is not possible to examine the effects of silencing specific pathways.

While acute hippocampal slices preserve the local tissue structure, many afferent connections are lost, causing strong activity alterations. In addition to the global deafferentation, it is possible to decrease synaptic input from specific pathways and evaluate reactive changes. However, whether homeostatic-like compensations after deafferentation can occur during the lifespan of acute slices, requires examination. In slices, CA3 pyramidal neurons display substantial firing and at least 20% of spontaneous EPSCs in CA1 neurons are due to CA3 intrinsic activity (Banerjee et al., 2013). Thus, it is reasonable to expect that this form of deafferentation could induce synaptic upregulation. In line with this possibility, a recent study analyzed evoked field excitatory postsynaptic potentials (fEPSP) in Schaffer collateral inputs to CA1 in slices that underwent CA3 removal, reporting a rapid (~3 h) increase in synaptic strength specifically in this pathway, compared to controls with intact CA3 (Dumas et al., 2018). However, the mechanism enabling these synaptic compensations require future evaluation.

Interestingly, fast (hours) adaptations to inactivity can be triggered by combining TTX and NMDAR antagonists (Sutton et al., 2006; Ibata et al., 2008) in cell cultures or acute slices, by a mechanism different from slow forms of HSP. To explore the nature of synaptic changes developed over the lifespan of slices that underwent CA3 dissection, we compared mEPSCs at a relatively early period (5-8 h) with those at a later interval (9-12 h). We observed distance-dependent compensations in mEPSC amplitude and frequency that were differentially affected by CaMKII inhibition. Our evidence supports that the mechanisms underlying these compensations resemble distinct known drug-induced forms of HSP.

## MATERIALS AND METHODS

Animal care and experimental procedures were approved by the Bio-Ethics Committee of the Facultad de Ciencias, Universidad de Chile, according to the Biosafety Policy Manual of the Fondo Nacional de Desarrollo Científico y Tecnológico (FONDECYT), Chile.

### Hippocampal slice preparation and preincubation

Male Sprague Dawley rats, 18-22 days old, were deeply anesthetized and decapitated. Transverse hippocampal slices (350 μm) were prepared in an ice-cold dissection solution containing (in mM) 125 NaCl, 2.6 KCl, 10 MgCl_2_, 0.5 CaCl_2_, 26 NaHCO_3_, 1.23 NaH2PO_4_, and 10 D-glucose (equilibrated with 95% O_2_ and 5% CO_2_), pH 7.3. CA3 was surgically removed and slices were transferred to interface chambers (tissue inserts, 8 μm) in an atmosphere saturated with 95% O_2_ and 5% CO_2_, at 30 °C. Each insert held 2-3 slices submerged in a drop of 200 μl artificial cerebrospinal fluid (ACSF) containing (in mM) 125 NaCl, 26 NaHCO_3_, 1 NaH2PO_4_, 2.6 KCl, 2 CaCl_2_, 1 MgCl_2_, and 10 D-glucose. After 1 h, the drop of ACSF was gently replaced by freshly-oxygenated solution (for Figs. 1 and 2). Slices were kept for 5-12 h in the chambers. For peptide preincubations, solution was replaced by oxygenated ACSF containing CN21 or SCR peptide (5 μM). After 2 h, slices were washed four times and maintained for at least 2 h before recordings.

**Figure 1.**
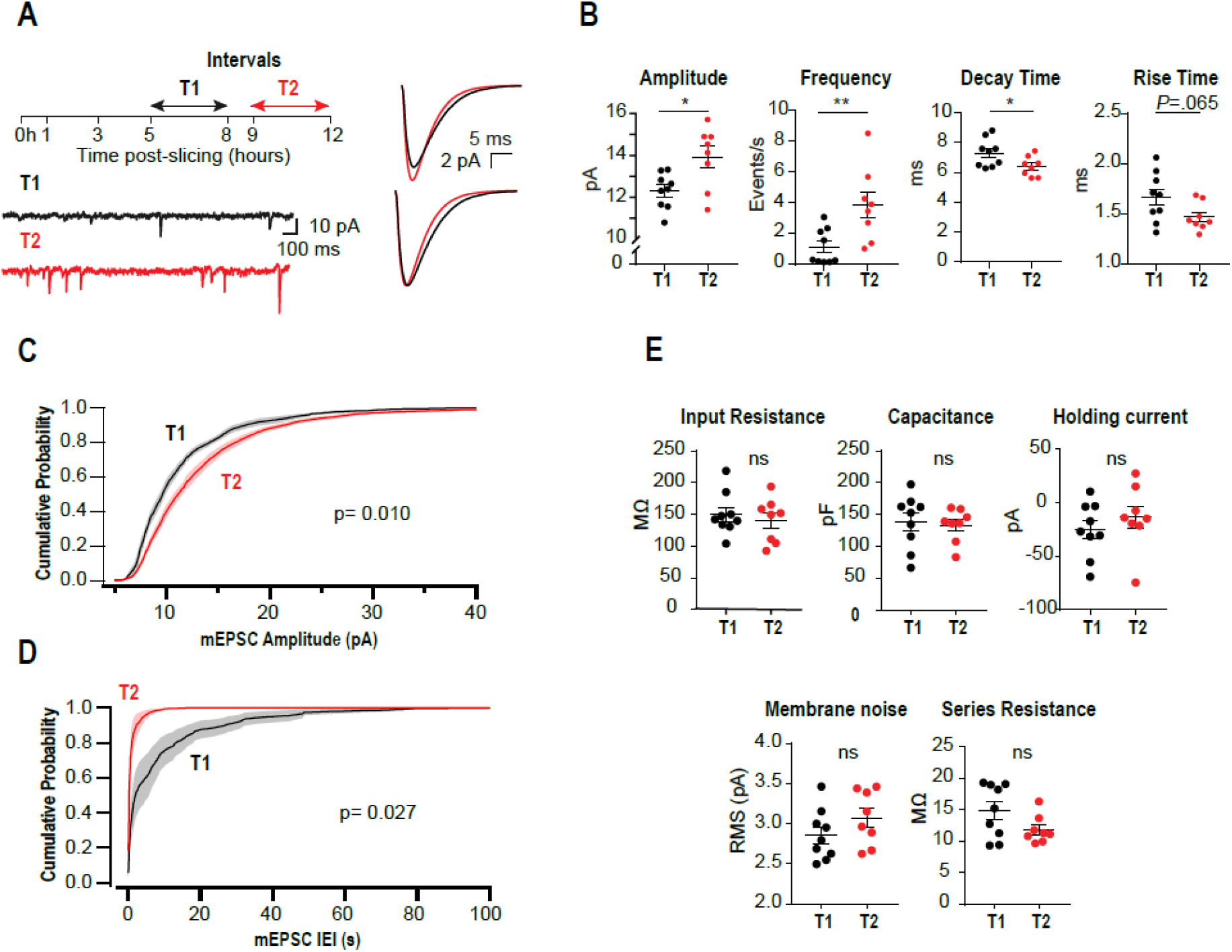
Upregulation of mEPSCs amplitude and frequency in acute slices lacking CA3. **(A)** Left, experimental design and representative somatic current recordings from two different neurons at early (T1) and late (T2) stages. Right, superimposed average mEPSCs waveforms (above) and rescaled traces (below). **(B)** Summary plots for mEPSCs amplitude, frequency, rise and decay times for all recorded neurons. Each point corresponds to the mean value from a single neuron. **(C)** Amplitude cumulative probability distributions of all mEPSC recorded at T1 or T2. **(D)** Interevent interval (IEI) distributions. Random permutation test. **(E)** Membrane properties and series resistance for the two cell populations (Unpaired t-test). N = 9 cells for T1 and N = 8 for T2.

**Fig. 2.**
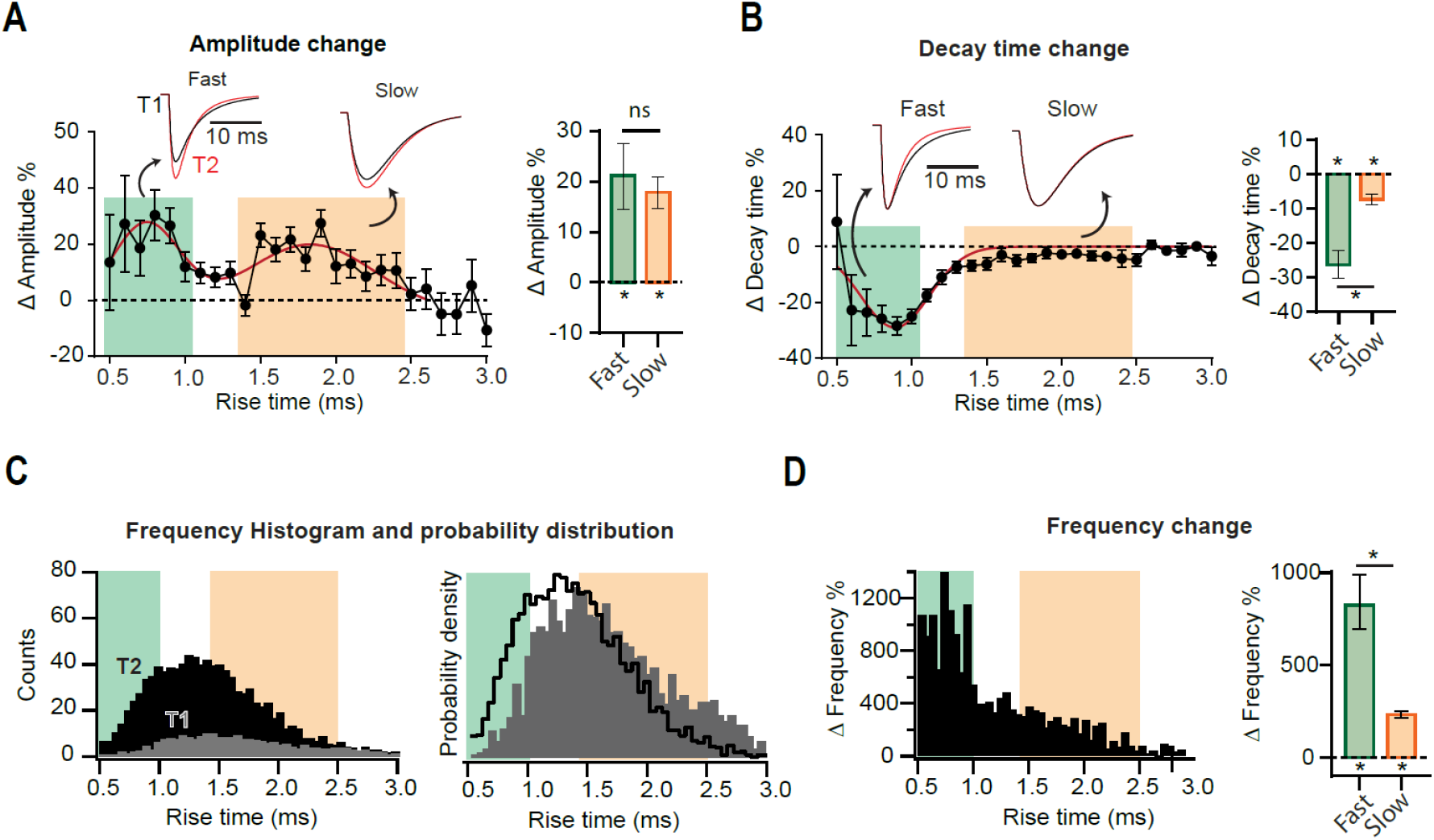
Heterogeneous synaptic adaptations in different groups of synapses. All mEPSCs recorded at a time interval (T1 or T2) were pooled together and sorted by rise time (***τ***_***r***_). **(A)** Left, distribution of mEPSC amplitude percent changes for ***τ***_***r***_-binned data (0.1 ms). The red curve corresponds to a bimodal fit (fast and slow-rising mEPSCs). Insets, superimposed average traces for T1 (black) and T2 (red), rise-time values indicated by the arrows. Data was separated into fast and slow rising groups using a 95% confidence interval obtained from the multi-peak fit. Right, mean amplitude percent increase for slow and fast-rising mEPSC. **(B)** Left, decay time (***τ***_***d***_) percent change distribution for ***τ***_***r***_-binned data. Right, decay time change summary plot for slow- and fast-rising mEPSC. **(C)** Left, superimposed frequency histograms for all events recorded at T1 or T2. Right, probability density curves, showing a relative increase in the incidence of fast-rising events. **(D)** Left, ***τ***_***r***_-binned frequency percent increases. Right, average frequency increase for fast and slow events. Bootstrap with Bonferroni correction (bar plots in A, B and F). N = 9 cells for T1 and N = 8 for T2. \**P*<0.05.

### Peptides

The CaMKIIN-derived peptide CN21 fused to the cell-permeable sequence tat (Vest et al., 2007; Sanhueza et al. 2011) and a scrambled control peptide, were obtained from Biomatik (Wilmington, Delaware, USA).

### Electrophysiological recordings

Slices were transferred to a submersion-type recording chamber mounted on an upright microscope (Nikon E600FN) and were continuously superfused (2–4 ml/min) with oxygenated ACSF. A total volume of 10 mL solution was recirculated using a two-way pump. Whole-cell patch-clamp recordings were performed using an EPC-10 amplifier (HEKA Elektronik, Reutlingen, Germany). Patch electrodes (2-5 MΩ electrodes) were pulled from borosilicate glass and the internal solution contained (in mM): 115 cesium methanesulfonate, 20 cesium chloride, 0.6 EGTA, 10 HEPES, 4 Na_2_-ATP, 0.4 Na-GTP, 10 Na_2_-phosphocreatine and 2.5 MgCl_2_ (293 mOsm, pH 7.25). mEPSCs were measured at −60 mV without junction potential (~12 mV) correction. Recordings were performed at 30 ± 1°C in the presence of 100 μM picrotoxin and 1 μM TTX.

### Statistics and data processing

Data analysis was done using GraphPad 7 (Prism) and custom-built algorithms designed in IgorPro 8 (Wavemetrics, Inc.). Equality of variance and normality were addressed using the Brown-Forsythe and the Shapiro-Wilk normality test, using an α value of 0.05. In Fig. 2 and Figs. 3F, L, error bars are the 95% confidence interval of the mean. In all other figures error bars are SEM.

**Figure 3.**
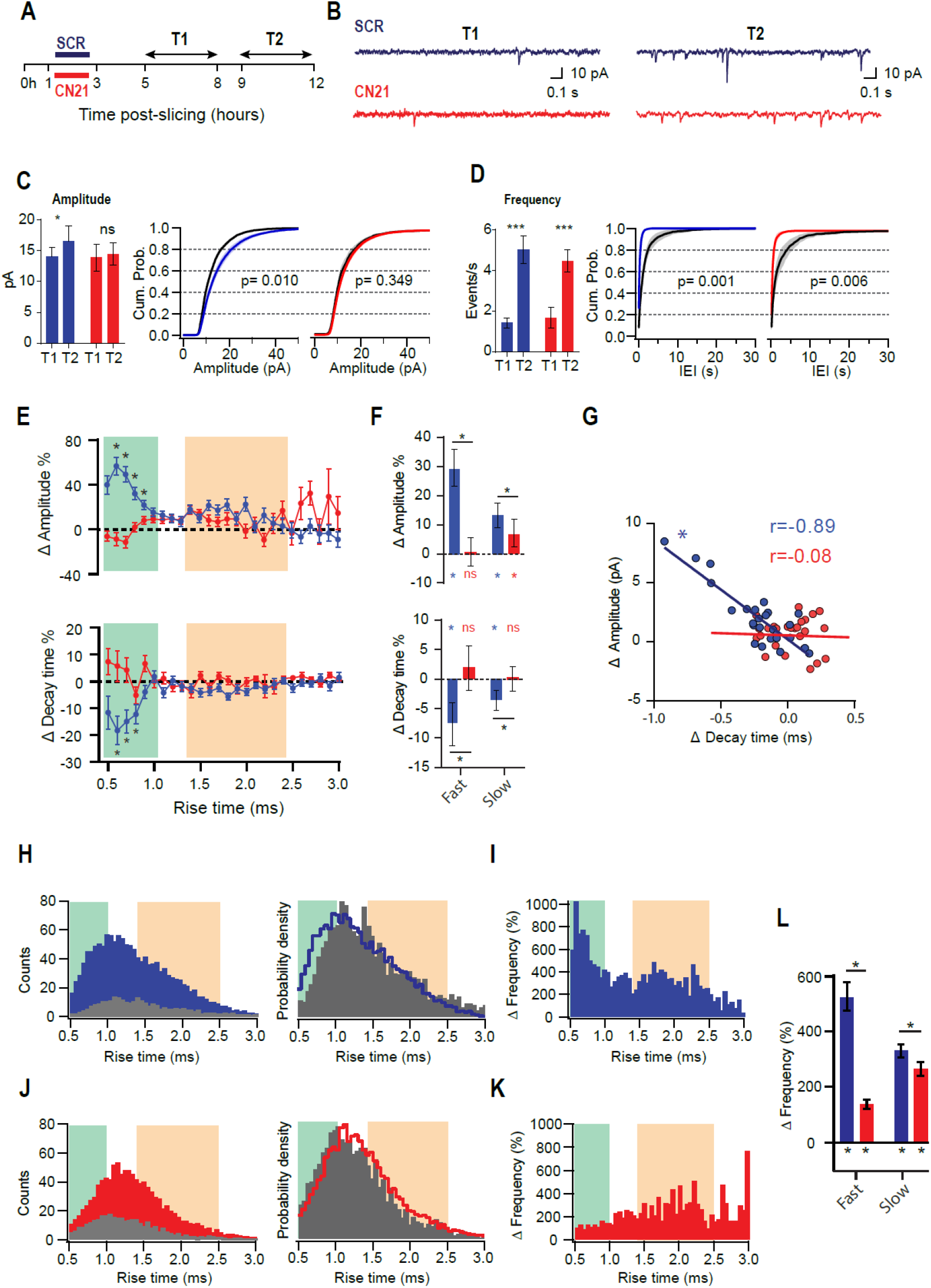
CaMKII is involved in different adaptations in acute slices. **(A)** Experimental design. Slices were preincubated for 2 h with CN21 or SCR peptide and drugs were washed out for 2 h (T1 and T2 are the same as for Figs. 1 and 2). **(B)** Representative current traces recorded at T1 and T2 in each condition **(C)** Mean mEPSCs amplitudes at T1 and T2 (left) and overlaid cumulative probability curves for CN21 (center) and SCR (right), including all mEPSCs recorded during each period. Black curves: distributions at T1. Two-way ANOVA with Bonferroni correction for bar plot, random permutations for cumulative probability plots. **(D)** Same as (C) for mEPSCs frequency and interevent interval (IEI). **(E)** Percent changes in amplitude and decay time for ***τ***_***r***_-binned data. Bootstrap with weighted P-values. **(F)** Average amplitude and decay time percent changes for slow and fast-rising groups. Bootstrap with Bonferroni correction. **(G)** Amplitude versus decay time raw changes for CN21 or SCR peptide. Each point corresponds to the average change for ***τ***_***r***_ bin. Pearson correlation. **(H)** Frequency histograms (left) for all events recorded at T1 or T2 and probability density curve (right), in SCR condition. **(I) *τ***_***r***_-binned frequency percent increase. (**J, K)** Same as (H-I), for CN21. **(L)** Average frequency increase for fast and slow groups, after SCR or CN21. Bootstrap with Bonferroni correction. \**P*<0.05. N = 8 cells for SCR-T1, 11 for SCR-T2, 12 for CN21-T1, and 11 for CN21-T2.

#### Random permutations test

Cumulative probability plots were compared by calculating the maximum absolute distance between the curves obtained after averaging distributions for all neurons recorded at T1 or T2. To evaluate statistical significance, surrogate data were produced by 10,000 random permutations of the cumulative plots.

#### Bootstrap with weighted p-value

To compare rise-time-binned data in Fig. 4, a p-value was calculated for each bin with the percentile bootstrap test. Family-wise error rate was corrected by multiplying each p-value by that of adjacent neighbors. This maintains family-wise error below 5% and preserves regional inference (Yang et al., 2001).

**Figure 4.**
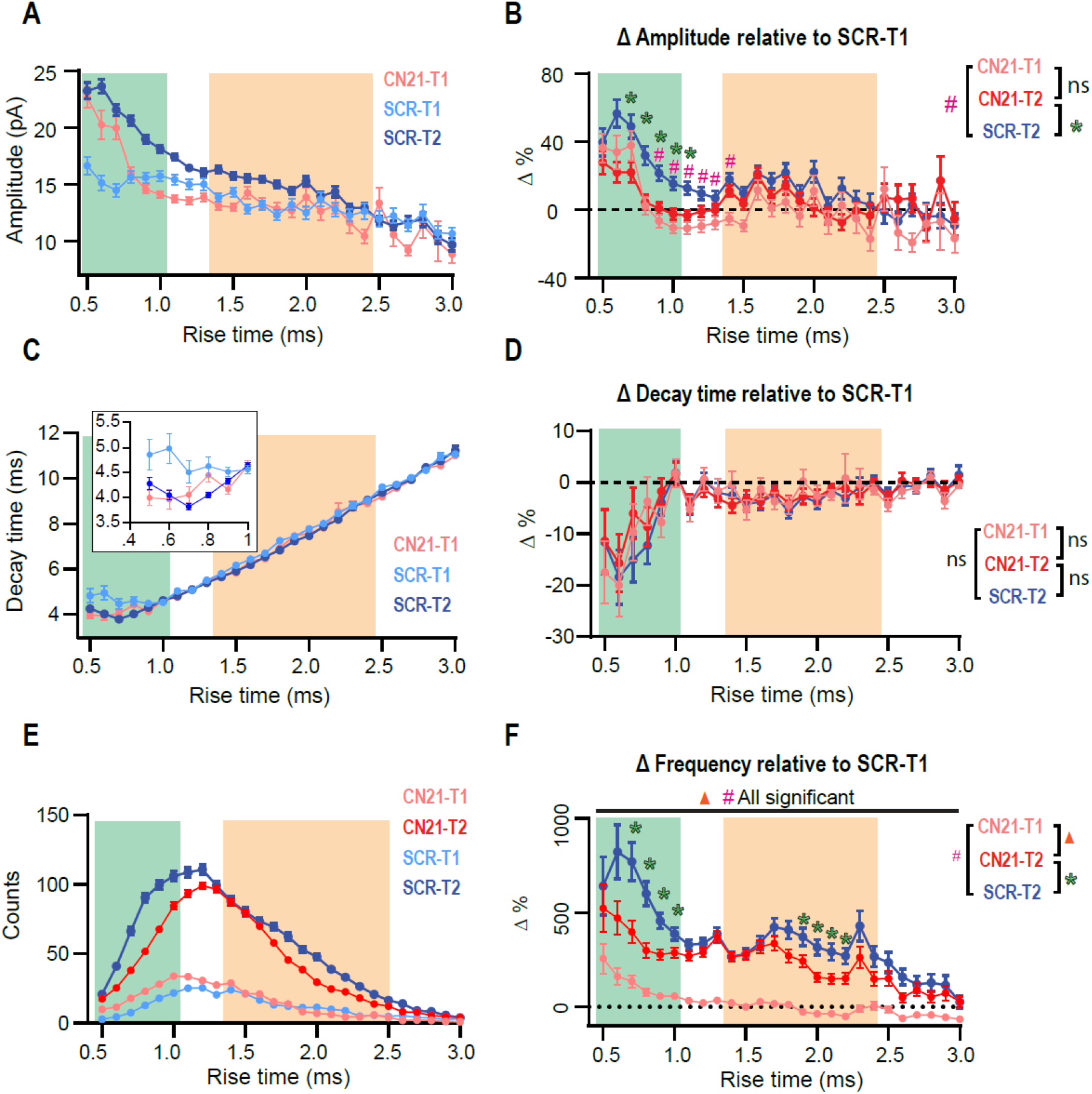
Early CaMKII inhibition abolishes or occludes homeostatic adaptations in different groups of synapses. **(A)** Overlaid ***τ***_***r***_-sorted mEPSC amplitude distributions for each temporal stage and condition. The curve for CN21 at T2 was omitted for simplicity. **(B)** Amplitude percent differences with respect to the SCR distribution at T1 **(C)** Same as (A), for decay time. Inset, detail for fast-rising group. **(D)** Same as (B), for decay time. **(E)** Superimposed event frequency distributions for each time interval and condition (same curves as in Fig. 3H, J; left). (F) Frequency increases with respect to SCR condition at T1. Bootstrap with weighted P-values (all panels). \**P*<0.05.

## RESULTS

To evaluate time-dependent synaptic adaptations in CA1 pyramidal neurons from acute slices lacking CA3, we recorded mEPSCs at two different intervals after preparation: T1 (5-8 h) and T2 (9-12 h) (Fig. 1A, left). Earlier than 5 h, mEPSCs frequency was too small to perform statistical analyses. Fig. 1A, right, displays the overlaid average mEPSC waveforms for all events recorded at T1 or T2, and their rescaled version (below) suggesting modifications in amplitude and kinetics. Accordingly, summary plots in Fig. 1B show a significant increase in amplitude and a decrease in decay time, each point corresponds to the mean value from one cell. Moreover, a strong increase in average frequency was demonstrated. Figs. 1C and D display amplitude and frequency cumulative probability distributions, respectively, for T1 and T2 (see summary data and statistics for Fig. 1 in Supplementary Table 1). These results are consistent with a compensatory process tending to recover transmission in CA1 neurons after the abrupt decrease in activity by slicing and CA3 dissection. Interestingly, in classical studies of HSP a decrease in ***τ***_***d***_ has been associated with the insertion of GluA2-lacking receptors displaying faster decay kinetics (Turrigiano, 2008; Lee, 2012) (Geiger et al., 1995).

To keep slices healthy during such long incubation times, we used interface chambers (Gibb and Edwards, 1994; Lipton et al., 1995). We checked for possible alterations in neuronal membrane properties after long-lasting storage, finding that passive properties, holding current, and membrane noise were similar for T1 and T2 (Fig. 1E). Moreover, there was no significant change in series resistance. Thus, changes in mEPSCs cannot be attributed to time-dependent modifications in these variables.

To explore possible differential adaptations along the dendritic tree, we sorted all recorded mEPSCs by their rise time (***τ***_***r***_), which is inversely correlated to the distance to the soma (Letzkus et al., 2006; Sjöström and Häusser, 2006). mEPSCs were grouped and averaged in ***τ***_***r***_ bins (0.1 ms) to then compute the changes between T1 and T2. This analysis revealed that percent increase in amplitude followed a bimodal distribution (Fig. 2A), possibly involving two separated synapse groups. A third population (***τ***_***r***_ > ~2.5 ms), presumably located most distally, did not display detectable changes. To quantify the average change in amplitude for each population, we established a quantitative criterium to classify as “fast” or “slow” the points in the distribution. Data was separated into two groups delimited by a 95% confidence interval, obtaining the following limits: fast population, ***τ***_***r***_ ≤ 1 ms (green) and slow population, 1.4 ms ≤ ***τ***_***r***_ ≤ 2.5 ms (pink). Events lying in the transition among the two groups (1.0 ms ≤ ***τ***_***r***_ ≤ 1.4 ms), cannot be classified as belonging to a specific group. The insets in Fig. 2A show superimposed average mEPSCs at T1 and T2, calculated from all events in each population. mEPSCs amplitude increase was significant for both fast and slow groups (Fig. 2A, right), with no statistical difference among them (see summary data and statistics for Fig. 2 in Supplementary Table 2). Comparing the decay time (***τ***_***d***_) as a function of ***τ***_***r***_, reveled evident changes for the fast-rising group, while the second population presented a very small (but significant) modification (Fig. 2B).

To examine whether the increase in frequency followed a particular pattern, we constructed histograms of event counts as a function of ***τ***_***r***_, at T1 and T2. Superimposed curves (Fig. 2C, left) demonstrated a strong and widespread increment in mEPSC occurrence. Both distributions display a single peak, however, the probability density distributions (Fig. 2C, right), revealed a shift to the left at T2. Therefore, while the occurrence of both fast and slow-rising mEPSCs strongly increased with time, it was more probable to observe faster events compared to slower ones at T2 than it was at T1. In line with this result, percent frequency increase was substantially higher in the lower ***τ***_***r***_ range (Fig. 2D). With regards to the slowest events (***τ***_***r***_ > 2.5 ms), the increase was negligible.

Overall, these observations are consistent with the development of synaptic upregulation mechanisms in distinct populations of synapses along CA1 neuron dendrites along slices lifespan. As Schaffer collaterals provide a major input to these cells, and CA3 removal was shown to increase evoked field potentials (Dumas et al., 2018), these changes could be in part due to this pathway-specific deafferentation. Interestingly, in the domain displaying the highest frequencies at T1, amplitude increases were lower, consistent with a homeostatic-like effect.

To address whether CaMKII mediates synaptic adaptations in our slices as it does in some known forms of HSP, we inhibited the holoenzyme using the CN21 peptide, a 21 amino-acid sequence derived from an endogenous protein, that specifically and efficiently inhibits CaMKII. After recovery slices were incubated for 2 h with cell-permeable, tat-fused CN21 (Fig. 3A) to inhibit CaMKII activity during the early post-denervation period. A second group of slices was treated with a scrambled control peptide (SCR). After washing out drugs for at least 2 hours, we conducted interleaved mEPSCs recordings for each condition at the same T1 and T2 intervals as for previous experiments. As shown in Fig. 3C (left), the increase in mEPSC amplitude also developed after SCR but was abolished by CN21, as calculated by averaging the mean values from each cell (see summary data and statistics for Fig. 3 in Supplementary Table 3). Moreover, after active peptide incubation the cumulative probability curves at T1 and T2 were not statistically different (Figs. 3C, right). In contrast, an increase in average frequency was detected in both conditions (Fig 3D).

Analysis of ***τ***_***r***_-sorted mEPSCs amplitude changes revealed that incubation with CN21 abolished upscaling of fast-rising mEPSCs and significantly reduced it for the slow-rising group (Figs. 3E, F; upper panels). In addition, CN21 prevented ***τ***_***d***_ decrease for both populations (lower panels). These observations support a role of CaMKII in mEPSCs upscaling and suggest that this regulation may involve the incorporation of GluA2-lacking receptors. Consistent with this possibility, we found a linear inverse correlation between amplitude increases and ***τ***_***d***_ decreases (Fig. 3G), suggesting a shared signaling pathway. This correlation was missing after CN21 incubation.

Examination of ***τ***_***r***_-sorted mEPSCs frequencies revealed that while an upregulation was observed after both SCR and CN21, the distribution was quite different (Fig. 3H-K). The strong increase in frequency of fast-rising events was decreased by CN21, falling below the level of the slow group (Fig. 3I, K, L). In addition, the increment in slow events incidence was also dampened after CN21. These results point to the existence of both CaMKII-dependent and independent mechanisms for frequency raise.

In summary, we detected CaMKII-related synaptic compensations developing hours after dissection. Interestingly, a closer examination of these reactive adjustments revealed further complexities consistent with the existence of different CaMKII-related mechanisms. We will next show that early kinase inhibition produced either suppression or occlusion of changes in separate synapse populations.

So far, we had focused on quantifying changes between two time periods (T1 and T2) in slices treated with either active or control peptides. However, Dumas et al (2018) reported that dissection of CA3 caused fEPSP upregulation after ~3 h, thus, some synaptic changes could be already in course at T1 in our slices. Therefore, we compared the amplitude distributions for SCR- and CN21-incubated slices at T1 (Fig. 4A, light blue and pink traces) and included the control curve at T2 for comparison (blue trace). At T1 the curves clearly differ for some ***τ***_***r***_ domains (see details on statistics in Supplementary Figure 1A). Intriguingly, after CN21, the mEPSCs displaying the fastest rising kinetics were already augmented at T1. On the other hand, a different population was lower compared to control. To visualize and quantify changes among conditions and time, we plotted the amplitude differences relative to the control at T1, now also including the curve for CN21 at T2 (Fig. 4B). Results suggest that CaMKII inhibition promoted the earlier development of amplitude compensations in part of the fast-rising population and in the slow group. Consistent with Fig. 3E, early and late curves from CN21-treated slices did not differ, but this may be due to two different phenomena: first, an occlusion effect for the fastest subpopulation and the slow group; second, a proper inhibition of the amplitude increase for an intermediate group. A similar analysis of decay times (Figs. 4C, D) also suggests a faster reorganization in AMPARs subunit composition after CN21.

Regarding frequency changes, Fig. 4E displays overlaid histograms (same as in Figs. 3H, J; left) and Fig. 4F, the percent differences relative to control at T1. Again, after CN21 the increase in the faster synaptic events in frequency appeared to start earlier (pink traces in Fig. 4F and Supplementary Figure 1C). However, the difference between pink and red curves in Fig. 4F confirms that frequency upregulation also included a global CaMKII-independent mechanism. Finally, comparing these curves with the control increment (Fig. 4F, blue curve) reveals that CN21 also inhibits a frequency increase depending on CaMKII and involving both fast and slow-rising events. Interestingly, like the amplitude increases, CaMKII-sensitive frequency changes were higher for synapses with lower incidence at T1 and were practically absent for the more abundant population, suggesting that the compensatory changes observed in our model of tissue denervation resemble homeostatic plasticity mechanisms involving CaMKII.

## DISCUSSION

We examined mEPSCs changes along the lifespan of acute slices that underwent CA3 removal. These compensations did not require pharmacological manipulations, were differentially expressed along dendrites, and involved distinct CaMKII-related regulations. As will be discussed, our results support that acute denervation can rapidly induce, in a single neuron, different synaptic adjustments resembling classical forms of adaptation to inactivity.

Average mEPSCs amplitude and frequency increased over early (T1, 5-8 h) and late (T2, 9-12 h) stages. As earlier than 5 h synaptic activity was very low, these changes are consistent with an ongoing inactivity-induced compensatory process. To investigate the underlying mechanisms, we sorted mEPSCs by their rise time (***τ***_***r***_), which depends on dendritic filtering and provides an estimate of the relative distance to the soma. mEPSCs presumably originating at separate regions of the dendritic tree followed nonuniform rules and the domains with higher activity levels at T1 developed lower amplitude and frequency increases, consistent with locally-regulated compensatory phenomena.

Mean amplitude increase was accompanied by a decrease in decay time (***τ***_***d***_), which is usually interpreted as a rise in the proportion of GluA2-lacking receptors, due to their faster deactivation/desensitization kinetics (Geiger et al., 1995). Moreover, amplitude and decay time changes displayed a linear inverse correlation, further suggesting a major contribution of this type of AMPARs to synaptic strength upregulation. Interestingly, several lines of evidence support a role for GluA2-lacking AMPARs in homeostatic upregulations after long-lasting neuronal activity silencing, including pharmacological manipulations, sensory deprivation, and inhibition of glutamate release at single synapses (Ju et al., 2004; Thiagarajan et al., 2005; Goel et al., 2006; Hou et al., 2008; Beique et al., 2011; Groth et al., 2011). A similar mechanism underlies the faster compensations triggered by simultaneous action potential and NMDAR inhibition (Sutton et al., 2006; Wang et al., 2011). Moreover, consistent with our observations, an increase in GluA1 content was detected by immunohistochemistry specifically in *stratum radiatum* 3 h after CA3 dissection, compared to slices preserving this region (Dumas et al., 2018).

We observed a global raise in mEPSC frequency, but increases were higher at the loci where amplitude changes occurred, consistent with both specific and unspecific frequency compensations. Some studies of adaptations to inactivity have reported frequency increases (Thiagarajan et al., 2005; Echegoyen et al., 2007; Groth et al., 2011) and may involve an enlargement of presynaptic terminals or an increase in vesicle pools and turnover (Bacci et al., 2001; Murthy et al., 2001). Interestingly, release probability can be homeostatically regulated by the local dendrite activity (Branco et al., 2008), consistent with a lower frequency increase for originally more active synapses. Finally, an overall increase in spine density and multiple-synapse boutons occur in CA1 neurons hours after slice preparation (Kirov et al., 1999). Moreover, drug-induced silencing triggered widespread spine increases in slices (Kirov and Harris, 1999). As in these studies CA3 was preserved, the cell-wide frequency increase observed here may be in part due to a general synapse upregulation triggered by the slicing procedure itself.

Different lines of evidence relate CaMKII with slow drug-induced adaptation to inactivity. Low Ca^2+^ levels preferentially activate CaMKIIβ, and knockdown of this subunit or general CaMKII pharmacological inhibition abolished mEPSC amplitude and frequency rise, supporting a specific role of ß in this form of HSP (Thiagarajan et al., 2002; Groth et al., 2011). Consistently, CaMKIIβ-dependent phosphorylation of the AMPAR regulatory protein stargazin is required for receptor incorporation to synapses (Louros et al., 2014). Moreover, phosphorylation of the guanylate-kinase-associated protein (GKAP) by CaMKIIβ boosts its synaptic accumulation and assembling of a scaffold complex with PSD-95 and Shank (Naisbitt et al., 1999; Shin et al., 2012; Rasmussen et al., 2017). However, the decrease in CaMKIIα activity during network silencing may also contribute to homeostatic upregulation. Normally, GKAP phosphorylation by CaMKIIα promotes its removal from synapses, limiting synaptic strength. Moreover, CaMKIIα-dependent proteasome recruitment causes degradation of GKAP and other scaffolds, destabilizing receptors and mimicking activity-induced downscaling (Bingol et al., 2010; Djakovic et al., 2012; Rasmussen et al., 2017). Therefore, while CaMKIIβ may trigger synaptic upregulation, CaMKIIα activity is expected to restrict transmission levels.

In consequence, it is plausible to propose that we are observing two different types of CaMKII-dependent fast homeostatic synaptic compensations. Early CN21 application blocked the changes requiring CaMKII activity, consistent with the known CaMKIIβ-mediated mechanism. On the other hand, CaMKII inhibition hastened other compensatory changes, in line with an alteration of the CaMKIIα-dependent mechanism limiting synaptic strength. If this interpretation is correct, the question arises on the factors determining which mechanism is expressed or prevails in different synapses.

It could be hypothesized that the effects observed here recapitulate, in part, the fast-developing HSP induced by TTX+APV (Sutton et al., 2006; Wang et al., 2011). These fast adaptations critically depend on CaMKIIα autophosphorylation and binding to NMDAR in cortical neurons (Wang et al., 2011), thus CN21 application could be mimicking NMDAR blockade. Moreover, mEPSC upregulation by TTX+APV involves locally-translated GluA2-lacking receptors, allowing rapid and specific adjustments (Sutton et al., 2006; Ju et al., 2004). Even individual synapses can autonomously compensate for neurotransmitter decreases (Ju et al., 2004; Hou et al., 2008; Beique et al., 2011), and Ca^2+^ entry through GluA2-lacking AMPARs can retrogradely increase vesicle turnover (Lindskog et al., 2010). The decay time shortening and differential changes along dendrites in our slices are consistent with these homeostatic mechanisms, and it could be hypothesized that a similar coordination among pre and postsynaptic modifications may underlie the coincident increase in amplitude and frequency observed here in groups of synapses. However, we also detected CN21-sensitive changes lacking a decay time reduction that may not rely on GluA2-lacking receptors. Further research is needed to assess these questions.

While mEPSC rise-time depends on synapse distance to the soma, other factors as dendrite diameter and branching may also affect this variable, and it is not straightforward to predict the relative position of synapses. While apical spine density in CA1 neurons is highest at the distal third of *stratum radiatum,* synapses are also abundant at basal dendrites, innervated by Schaffer collaterals as well (Megı́as et al., 2001). These questions should be addressed in future studies.

Interestingly, CN21 peptide derives from the CaMKIIN protein, a specific endogenous inhibitor of CaMKII (Chang et al., 1998; Vest et al., 2007), whose function is not well understood, but has been associated with Hebbian synaptic plasticity and memory (Sanhueza et al., 2011; Gouet et al., 2012; Vigil et al., 2017). Notably, the CaMKIIN gene is rapidly regulated after LTP induction (Astudillo et al., 2020). Therefore, our results may further contribute to understand possible CaMKIIN-dependent regulatory processes.

To the best of our knowledge, denervation-induced fast synaptic changes following similar rules than known forms of HSP, had not been demonstrated. Moreover, the simultaneous occurrence in a neuron of different homeostatic processes has not been reported. HSP studies usually compare control and test conditions at a similar time stage, in contrast to our work that investigates the development of changes over time. The study of ongoing adaptations occurring in slices may contribute to unraveling rapid pathway-specific mechanisms and understanding reactive phenomena following brain injury.

Finally, researchers should be aware that synaptic compensations can occur over the lifespan of slices, possibly affecting results interpretation.

## Supporting information

Summary data and statistical tests

## AUTHOR CONTRIBUTIONS

P.V. performed the experiments and analysis, actively participated in the interpretation of results, and wrote the first draft of the manuscript. G.P. and J.V. contributed to experiments and results discussion. M.S. designed research and wrote the final version of the manuscript.

## ACKNOWLEDGEMENTS

This work was supported by the Fondo Nacional de Desarrollo Científico y Tecnológico, FONDECYT grant No. 1140700 (MS).

## AVAILABILITY OF SUPPORTING DATA

All the data are available upon request.

