## Supplementary material for "Heterogeneous CaMKII-dependent synaptic compensations in CA1 pyramidal neurons from acute slices with dissected CA3": Summary data and statistical tests

**Table 1. Related to Fig 1B,3.** Statistical test: Unpaired t-test.

| Data | T1 | T2 | T1 vs T2(p) |
|---|---|---|---|
| Amplitude (pA): | 12.31± 0.28 | 13.92± 0.52 | 0.013 |
| Frequency (events/s): | 1.103± 0.39 | 3.845± 0.86 | 0.008 |
| Decay time (ms): | 7.32± 0.31 | 6.42± 0.23 | 0.039 |
| Rise time (ms): | 1.66± 0.08 | 1.47± 0.05 | 0.065 |
| Input Resistance (MΩ): | 150.4± 11.08 | 141.2± 12.24 | 0.582 |
| Capacitance (pF): | 138.5± 14.11 | 133.3± 9.23 | 0.766 |
| Series Resistance (MΩ): | 14.83± 1.40 | 11.81± 0.77 | 0.091 |
| Holding current (pA): | -25.28± 8.51 | -13.66± 10.67 | 0.403 |
| Membrane noise (pA): | 2.858± 0.11 | 3.073± 0.12 | 0.196 |

**Table 2. Related to Fig 2A-C.** Statistical test: Percentile Bootstrap with Bonferroni correction.

| Data | Fast | Fast ≠ 0 (p) | Slow | Slow ≠ 0 (p) | Fast vs Slow (p) |
|---|---|---|---|---|---|
| Amplitude Δ%: | 21.16±3.50 | <0.0006 | 17.81 ± 1.53 | <0.0006 | >0.9999 |
| Rise time Δ%: | -26.42 ± 1.99 | <0.0006 | -7.30 ± 0.74 | <0.0006 | <0.0006 |
| Frequency Δ%: | 832.60 ± 67.20 | <0.0006 | 230.98 ± 10.03 | <0.0006 | <0.0006 |

**Table 3. Related to Fig 3C, D.** Statistical test: Two-way ANOVA with Bonferroni correction.

| | SCR | | | CN21 | | |
|---|---|---|---|---|---|---|
| Data | T1 | T2 | T1 vs T2 (p) | T1 | T2 | T1 vs T2 (p) |
| Amplitude (pA): | 14.05 ± 0.44 | 16.57 ± 0.87 | 0.0175 | 13.86±0.79 | 14.45±0.52 | >0.9999 |
| Frequency (events/s): | 1.43 ± 0.24 | 5.011 ± 0.68 | <0.0001 | 1.67 ± 0.55 | 4.48 ± 0.55 | 0.0009 |

**Table 3. Related to Fig 3F, L.** Statistical test: Percentile Bootstrap with Bonferroni correction.

| Data | | SCR | SCR ≠ 0 (p) | CN21 | CN21≠ 0 (p) | SCR vs CN21 (p) |
|---|---|---|---|---|---|---|
| Amplitude Δ%: | Fast | 29.26 ± 2.42 | <0.0012 | 0.82 ± 1.90 | 1 | <0.0012 |
| | Slow | 13.20 ± 1.37 | <0.0012 | 6.73 ± 1.62 | <0.0012 | 0.014 |
| Rise time Δ%: | Fast | -7.51 ± 1.42 | <0.0012 | 1.84 ± 1.48 | 1 | <0.0012 |
| | Slow | -3.59 ± 0.55 | <0.0012 | -0.15 ± 0.7 | 1 | <0.0012 |
| Frequency Δ%: | Fast | 521.86 ± 21.00 | <0.0012 | 132.41± 6.54 | <0.0012 | <0.0012 |
| | Slow | 330.87 ± 9.26 | <0.0012 | 265.49 ± 10.33 | <0.0012 | <0.0012 |
