## Supplementary Figure 1 for "Heterogeneous CaMKII-dependent synaptic compensations in CA1 pyramidal neurons from acute slices with dissected CA3"

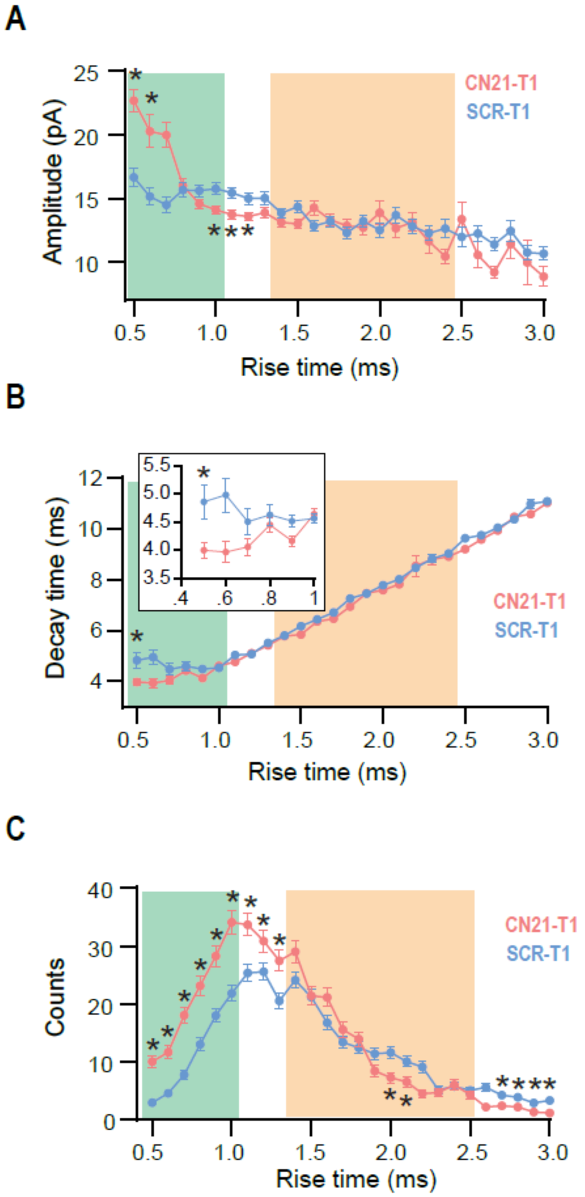


**Supplementary Figure 1. Synaptic modifications detected at T1 upon early CaMKII inhibition (A)** Overlaid $\tau_r$-sorted mEPSC amplitude distributions at T1, revealing that after CN21, transmission was already increased for the faster rising group and diminished for an intermediate population. (B) Same as (A), for decay time. Inset, detail of the curves for the faster-rising group, unraveling an earlier $\tau_r$ decrease after CN21 in this group. **(C)**

Comparison of event frequency distributions at T1 also shows an incipient upregulation of faster-rising events incidence after CN21.
